## Supplemental Figures for "Heterozygosity for neurodevelopmental disorder-associated *TRIO* variants yields distinct deficits in behavior, neuronal development, and synaptic transmission in mice"

### SUPPLEMENTAL FIGURES AND FIGURE LGENDS

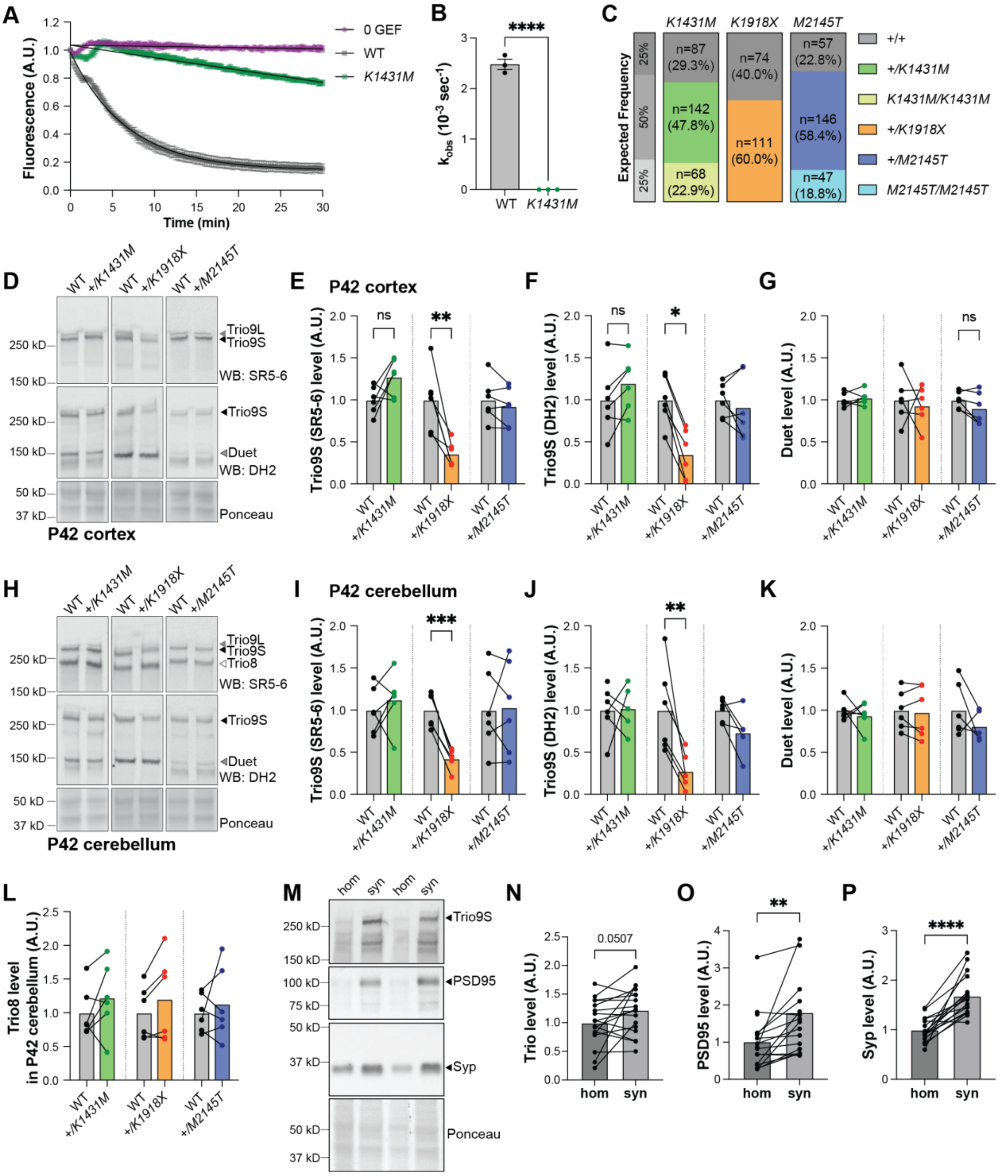


**Supplemental Fig. 1. Trio *+/K1918X* but not *+/K1431M* or *+/M2145T* mice have reduced levels of Trio protein in the brain.**

**(A)** *In vitro* GEF assay showing Rac1:Bodipy-FL-GDP exchange via fluorescence decay over time, with background subtracted and fitted with exponential curves. *K1431M* has impaired GEF activity compared to WT at equimolar amounts (500 nM) of TRIO GEF1 protein (n=3 replicates). **(B)** *K1431M* significantly decreases the initial rate of nucleotide exchange compared to WT ($K_{obs}$= 0.0006 ± 0.0002 x$10^{-3}$ $s^{-1}$ vs WT $K_{obs}$= 2.5 ± 0.1 x$10^{-3}$ $s^{-1}$; p<0.0001, n=3). **(C)** Number and frequency of progeny from heterozygote intercrosses. *+/K1431M x +/K1431M* crosses produced litters in the expected Mendelian frequencies of

25% +/+: 50% +/variant: 25% variant/variant (Chi-square test, two-tailed p=0.2231). *+/K1918X* (Chi-square test, two-tailed p<0.0001) intercrosses did not yield homozygote variants (binomial test, two-tailed p=0.0610). *+/M2145T x +/M2145T* crosses yielded slightly more heterozygotes than expected (Chi-square test, two-tailed p=0.0140). **(D,H)** Representative immunoblots for Trio isoforms in the cortex **(D)** and cerebellum **(H)** of P42 male *Trio* heterozygous variant and paired WT littermate mice (used antibodies noted in parentheses). **(E-G,I-L)** Quantification of Trio isoform levels in immunoblots of the cortex **(E-G)** and cerebellum **(I-L)**. Significant decreases in Trio9 levels were found only in *+/K1918X* cortex (in E: 0.3590 ±0.06005 vs WT 1.000 ±0.1541, p=0.0036; in F: 0.3513±0.1191 vs WT 1.000 ±0.1165, p=0.0207) and cerebellum (in I: 0.4239 ±0.04858 vs WT 1.000 ±0.08361, p=0.0003; in J: 0.2753 ±0.08413 vs WT 1.000 ±0.2118, p=0.0059, n=6 mice per group). No significant changes in Trio levels were observed in *+/K1431M* or *+/M2145T* mouse brains. Trio8 levels in the cerebellum for all *Trio* heterozygous variant mice were unchanged from WT littermates. Ratio paired t-tests identified differences from the WT mean (n=6 mice per group). **(M)** Representative immunoblots showing enrichment of Trio, PSD95, and synaptophysin (Syp) in synaptosomes (syn) compared to 40 µm-filtered total homogenate (hom) from P42 WT mouse cortex. **(N-P)** Quantification of Trio9S, PSD95, and Syp in immunoblots from P42 mouse synaptosomes versus total homogenate (n=18 mice). All data are presented as mean ± SEM; significance tested by Paired t-tests unless specified otherwise ($^{ns}$p<0.1, *p<0.05,**p<0.01, ****p<0.0001).

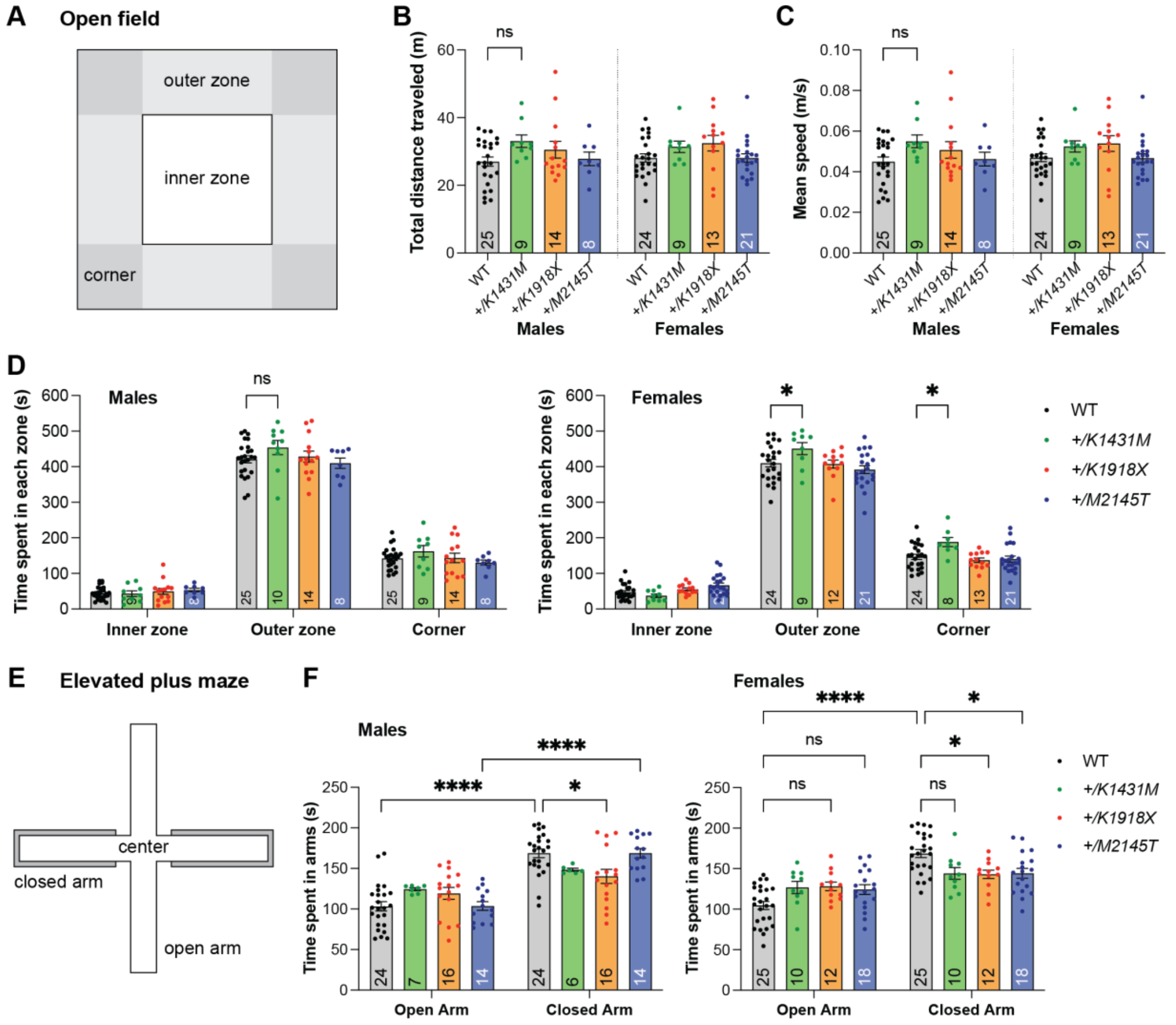


**Supplemental Fig. 2. Heterozygosity for distinct Trio variants differentially impact anxiety-like behaviors.**

**(A)** Schematic diagram of the open field test. **(B-F)** Total distance and time in each zone were tracked automatically over 10 minutes per mouse (n of mouse per group are shown inside the bar). **(B-C, D left)** *Trio* variant male mice did not show significant differences in the open field test compared to WT. **(D right)** *+/K1431M* females spent significantly more time in the outer and corner zones of the open field relative to WT females (outer zone: *+/K1431M* 450.622 ± 17.117 s, vs WT 409.774 ± 10.664 s zone, p=0.0157) (corner zone: *+/K1431M* 188.188 ± 12.876 s, vs WT 146.550 ± 7.251 s, p=0.0192). **(E)** Schematic diagram of the elevated plus maze. **(F)** WT mice of both sexes and *+/M2145T* males preferred to spend more time in the closed arms than in the open arms of the elevated plus maze; *+/K1431M* and *+/K1918X* mice of both sexes and *+/M2145T* females in time spent in open versus closed arms in the elevated plus maze test. Females of all *Trio* heterozygote genotypes as well as *+/K1918X* males displayed decreased time in the closed arm relative to WT mice, and exhibited trends towards increased time in the open arms relative to WTAll data are presented as mean ± SEM. Two-way ANOVA with post-hoc Bonferroni MC test identified differences from WT ($^{ns}$p<0.1, *p<0.05, ****p<0.0001). Numbers of mice quantified per group are annotated inside the bars.

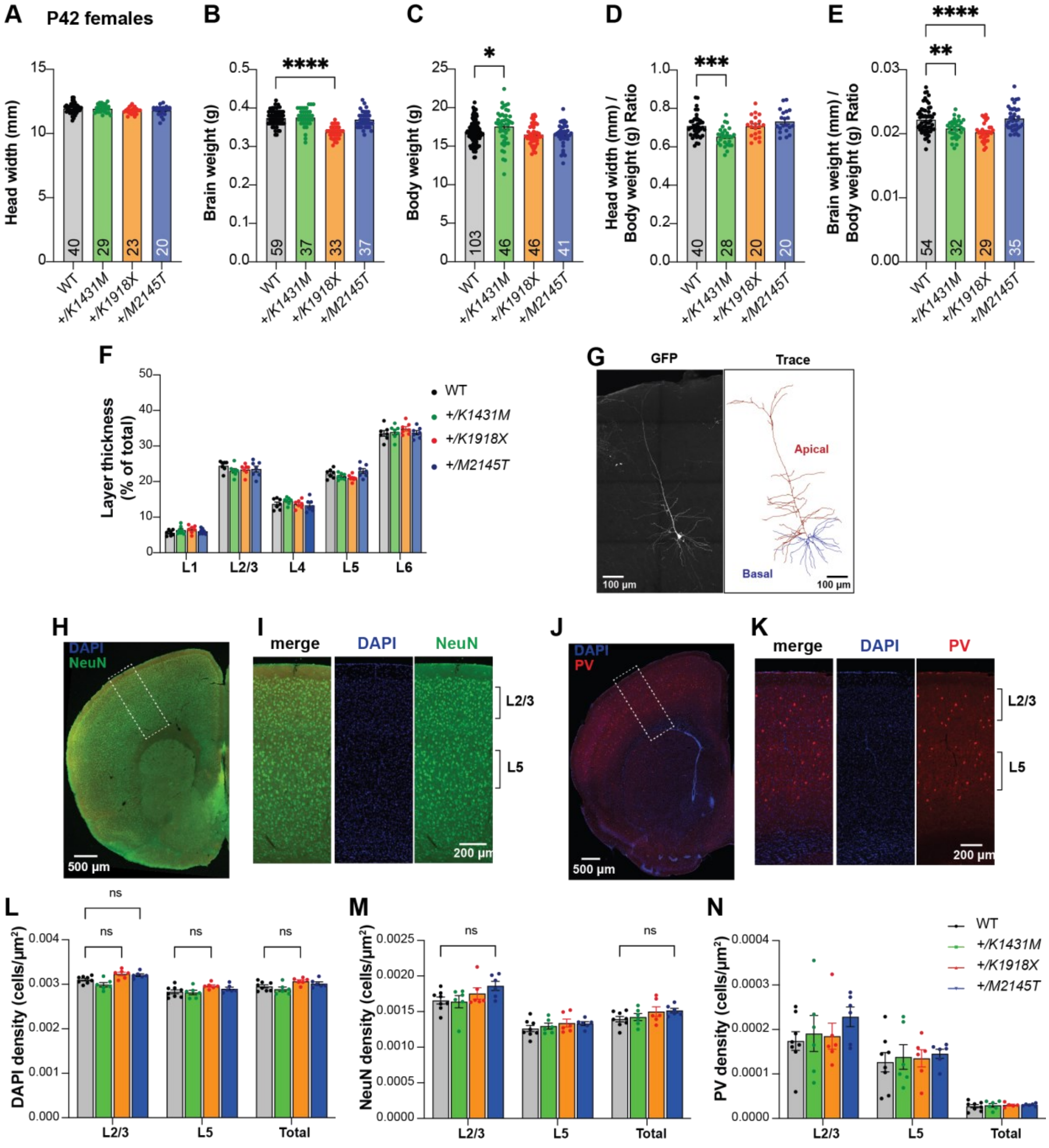


**Supplemental Fig. 3. Heterozygous *Trio* variant mice show mild alterations in cortical organization.**

**(A)** Ear-to-ear head width is unchanged from WT for all female heterozygous *Trio* variant mice at P42. **(B)** Brain weight is significantly reduced by 9.9% in P42 *+/K1918X* female mice compared to WT. **(C)** Body weight is increased by 4.2% in P42 *+/K1431M* female mice **(D)** Head widths normalized to body weight of P42 *+/K1431M* female mice were reduced by 7.9% compared to WT mice. Head width-to-body weight ratios were calculated per individual mouse, with mouse number per group annotated within the bar. **(E)** Brain weights normalized to body weight of P42 *+/K1431M* and *+/K1918X* female mice were reduced 6.2% and 8.6%, respectively, compared to WT mice. Brain-to-body weight ratios were calculated per individual mouse, with mouse number per group annotated within the bar. **(F)** Thickness of individual cortical layers expressed as a percentage of total cortical thickness. No differences were observed in *Trio* variant mice compared to WT. For (G-H), two-way ANOVA with post-hoc Bonferroni MC test identified differences from WT (**p<0.01; n=7 mice per group). **(G)** Representative maximum projection fluorescence image and corresponding dendritic arbor reconstruction of a motor cortex Layer 5 pyramidal neuron (M1 L5 PN) from a P42 *Thy1-GFP* mouse. **(H-K)** Representative fluorescent image of a 30 μm coronal slice from P42 WT brain, immunostained for NeuN and PV. Regions of motor cortex as outlined by the white dotted box in (I) and (K) are magnified in (J) and (L), resp. **(L-N)** Density of DAPI+, NeuN+, and PV+ cells did not significantly differ between *Trio* variants and WT in the total M1 region quantified or in cortical layers 2/3 and 5, though there were trends towards increased DAPI+ cell density in *+/K1918X* and increased L2/3 NeuN+ cell density in *+/M2145T* P42 male mice relative to WT. All data are shown as mean ± SEM; significance tested by one- way or two-way ANOVA with post-hoc Bonferroni MC test identified differences from WT (ns: $0.05<p<0.1$; n=6 mice for *Trio* variants, n=8 WT mice; 3 slices per mouse were analyzed). All data are presented as mean ± SEM.

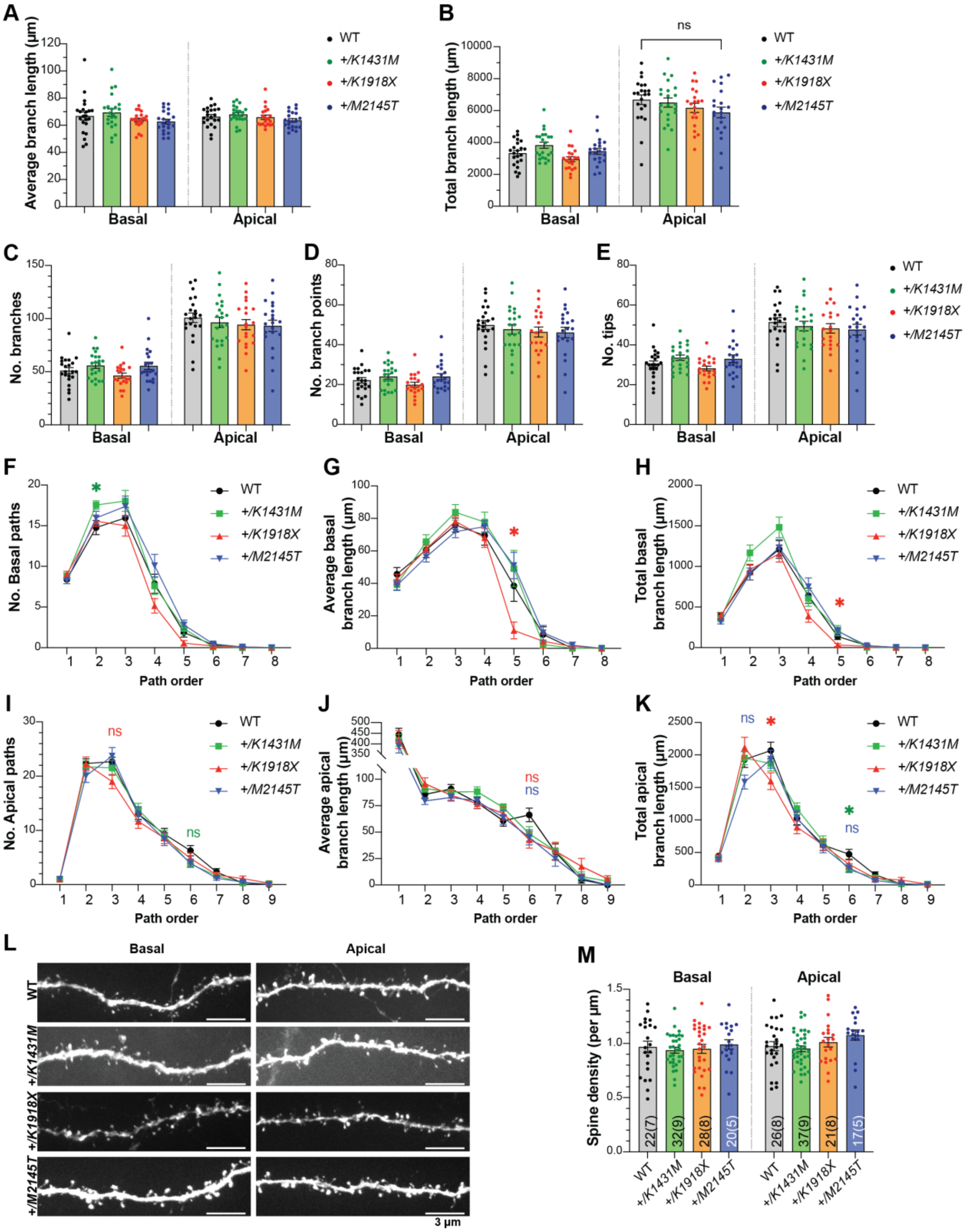

**Supplemental Fig. 4. Additional measurements of dendrites from M1 L5 PN reconstructions show modest order-dependent changes in *Trio* variant mice.**

**(A-E)** Overall measurements of dendritic reconstructions showed that average dendrite length (A), sum total dendrite length (B), number of branches (C), number of branch points (D), and number of tips (E) of *Trio* variant M1 L5 PNs were not different from WT. (n= same as in main Fig. 2J-M). **(F-K)** Path length analysis showed dendrite order-dependent changes in *+/K1431M* and *+/K1918X* number of branches (F,I), average dendrite length (G,J), sum total dendrite length (H,K) for basal (F-H) and apical (I-K) dendrites compared to WT.

*+/K1918X* PNs exhibited reduced average and total dendrite length at higher-order basal dendrites (in 5th-order basal dendrites: average branch length 11.015 ±5.176 um vs WT 38.51 ±9.565 um, p=0.0491; total dendrite length 31.593 ±18.700 um vs WT 137.550 ±38.407 um, p=0.0491) and reduced sum total branch length in mid-order apical dendrites (in tertiary apical dendrites, 1596.476 ±129.408 vs WT 2068.017 ±127.918 um, p=0.0354). *+/K1431* PNs had increased proximal basal dendrite numbers (in secondary basal dendrites, 17.545 ±0.513 um vs WT 14.783 ±0.866 um, p=0.0256). Two-way ANOVA with post-hoc Bonferroni MC test identified differences from WT **(L)** Representative maximum projection fluorescence images of basal and apical dendrite segments from M1 L5 PNs of P42 *Trio* variant mice. **(M)** Dendritic spine density on proximal apical and secondary basal dendrites are unchanged in *Trio* variant mice compared to WT (used 2-5 dendrites per neuron/1-2 neurons per mouse). Numbers of dendrites quantified per group are annotated inside the bar (number of neurons in parentheses). All data show mean ±SEM; significance tested using one-way or two-way ANOVA as appropriate with post-hoc Bonferroni MC test identified differences from WT ($^{ns}p<0.1$, *p<0.05, **p<0.01; n=same as in main Fig. 2J-M).

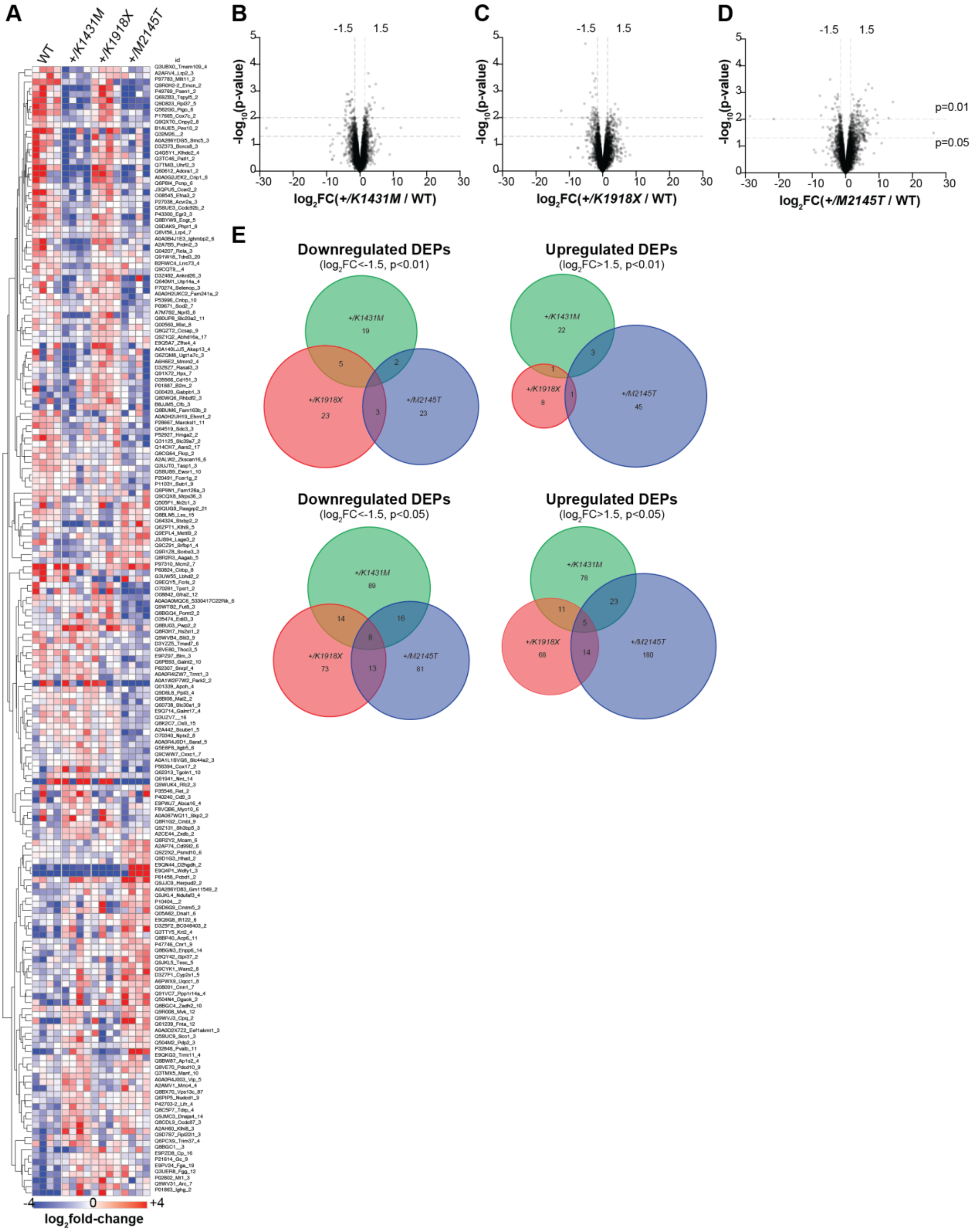


**Supplemental Fig. 5. Mass spectrometry-based proteomics reveals molecular changes in the brains of *Trio* variant mice compared to WT mice.**
**(A)** Heatmap of select protein abundances in P21 cortex of WT and Trio variant mice shows genotype differences. Each row is a protein, each column a mouse (n=4 mice per genotype). Displayed are 184/7362 proteins with significant differences ($p<0.01$, t-test). Red = upregulated, blue = downregulated, sorted by k-means clustering. Complete list of proteins shown in **Supp. Table 1**. **(B-D)** Volcano plots of differentially expressed proteins (DEPs) identified by proteomics for P21 *+/K1431M* (B), *+/K1918X* (C), and *+/M2145T* (D) mice,

expressed as $\log_2$(fold-change, FC) relative to WT mice (from n=4 mice per genotype). DEPs that are increased in the mutant compared to WT have a positive $\log_2$FC, while DEPs that are decreased in the mutant compared to WT have a negative $\log_2$FC. Dotted lines on y-axis indicate cutoffs of $p<0.01$ and $p<0.05$; dotted lines on x-axis indicate cutoffs of $\log_2FC<-1.5$ (downregulated DEPs) or $\log_2FC>1.5$ (upregulated DEPs). **(E)** Venn diagrams show little overlap in the up- and down-regulated DEPs between *Trio* variant mice, using DEP cut-off values of $\log_2FC>1.5$ (upregulated DEPs) or $\log_2FC<-1.5$ (downregulated DEPs), and $p<0.01$ or $p<0.05$.

### SUPPLEMENTAL TABLES

**Supp. Table 1. (Supp Fig.5 – Source data 1)** Proteome of P21 cortex from *Trio* WT, *+/K1431M*, *+/K1918X*, and *+/M2145T* mice.
